# Transcriptional and immunological analysis of the putative outer membrane protein and vaccine candidate TprL of *Treponema pallidum*

**DOI:** 10.1101/2020.09.28.316414

**Authors:** Austin M. Haynes, Mark Fernandez, Emily Romeis, Oriol Mitjà, Kelika A. Konda, Silver K. Vargas, Maria Eguiluz, Carlos F. Caceres, Jeffrey D. Klausner, Lorenzo Giacani

## Abstract

**Background:** An effective syphilis vaccine should elicit antibodies to *Treponema pallidum* subsp. *pallidum* (*T. p. pallidum*) surface antigens to induce pathogen clearance through opsonophagocytosis. Although the combination of bioinformatics, structural, and functional analyses of *T. p. pallidum* genes to identify putative outer membrane proteins (OMPs) resulted in a list of potential vaccine candidates, still very little is known about whether and how transcription of these genes is regulated during infection. This knowledge gap is a limitation to vaccine design, as immunity generated to an antigen that can be down-regulated or even silenced at the transcriptional level without affecting virulence would not induce clearance of the pathogen, hence allowing disease progression.

**Principal findings:** We report here that *tp1031*, the *T. p. pallidum* gene encoding the putative OMP and vaccine candidate TprL is differentially expressed in several *T. p. pallidum* strains, suggesting transcriptional regulation. Experimental identification of the *tprL* transcriptional start site revealed that a homopolymeric G sequence of varying length resides within the *tprL* promoter and that its length affects promoter activity compatible with phase variation. Conversely, in the closely related pathogen *T. p.* subsp. *pertenue*, the agent of yaws, where a naturally-occurring deletion has eliminated the *tprL* promoter region, elements necessary for protein synthesis, and part of the gene ORF, *tprL* transcription level are negligible compared to *T. p. pallidum* strains. Accordingly, the humoral response to TprL is absent in yaws-infected laboratory animals and patients compared to syphilis-infected subjects.

**Conclusion:** The ability of *T. p. pallidum* to stochastically vary *tprL* expression should be considered in any vaccine development effort that includes this antigen. The role of phase variation in contributing to *T. p. pallidum* antigenic diversity should be further studied.

**Author Summary:** Syphilis is still an endemic disease in many low- and middle-income countries and has been resurgent in high-income nations for almost two decades now. In endemic areas, syphilis still causes significant morbidity and mortality in patients, particularly when its causative agent, the bacterium *Treponema pallidum* subsp*. pallidum* is transmitted to the fetus during pregnancy. Although there are significant ongoing efforts to identify an effective syphilis vaccine to bring into clinical trials within the decade in the U.S., such efforts are partially hindered by the lack of knowledge on transcriptional regulation of many genes encoding vaccine candidates. Here, we start addressing this knowledge gap for the putative outer membrane protein (OMP) and vaccine candidates TprL, encoded by the *tp1031* gene. As we previously reported for other putative OMP-encoding genes of the syphilis agent, *tprL* transcription level appears to be affected by the length of a homopolymeric sequence of guanosines (Gs) located within the gene promoter. This is a mechanism known as phase variation and often involved in altering the surface antigenic profile of a bacterial pathogen to facilitate immune evasion and/or adaptation to the host milieu.

## Introduction

Syphilis is a chronic sexually transmitted infection that despite being relatively easy to prevent, diagnose, and treat, still represents a burden for public health as it causes significant morbidity and mortality worldwide. The World Health Organization estimates that syphilis global prevalence and incidence range between 18 to 36 million cases and between 5.6 to 11 million new cases every year, respectively [1, 2]. Although the majority of those cases occur in low- and middle-income countries where the disease is endemic, syphilis rates have also been steadily increasing for two decades in high-income countries, where men who have sex with men (MSM) and HIV-infected populations are affected [3–8]. In the US, for example, the rate of early syphilis in 2018 was 10.8 cases per 100,000 population, which represents a 414% increase compared to the 2.1 cases per 100,000 population reported in 2000 [3]. In absence of treatment, syphilis might progress to affect patients’ cardiovascular and central nervous systems, potentially leading to aortic aneurism, stroke, hearing or visual loss, dementia, and paralysis [9]. Furthermore, mother-to-child transmission of the infection during pregnancy accounts for up to 50% of stillbirths in sub-Saharan Africa and a high proportion of perinatal morbidity and mortality cases [10]. Additionally, in the US, the recent syphilis rate increase in women of reproductive age led to an increase in congenital syphilis cases from 362 cases in 2013 to 1,306 in 2018 [3]. Lastly, evidence that syphilis causes an approximate 5-fold increase in the likelihood of HIV transmission and acquisition [11] further highlights the threat posed by this disease to global health.

Overall, syphilis epidemiology supports the necessity for an effective vaccine to help disease control. Ongoing vaccine development efforts aim to elicit opsonic antibodies that target conserved surface epitopes of putative outer membrane proteins (OMPs) of the syphilis agent, the spirochete bacterium *Treponema pallidum* subsp. *pallidum* (*T. p. pallidum*), and confer sterilizing immunity by promoting opsonophagocytosis of *T. p. pallidum* by IFNγ-activated macrophages [12, 13]. That strategy, however, finds an obstacle in our limited knowledge of how this spirochete controls transcription of genes encoding vaccine candidates. Such knowledge is however pivotal to devising an effective vaccine, as antibodies generated against an antigen whose expression can be downregulated or even abrogated without affecting pathogen virulence or viability would be ineffective in clearing organisms not expressing the target. The only published high-throughput study that investigated the *T. p. pallidum* transcriptome used microarrays to provide a snapshot of the level of expression of OMP-encoding *T. p. pallidum* genes in treponemes harvested at peak orchitis from a rabbit infected with the Nichols strain, but could not address the topic of gene regulation [14]. Our past studies, on the other end, although limited to a subset of OMP-encoding genes, have suggested that transcription of several of those genes can be affected by a homopolymeric tract of guanosines (poly-G) of varying length located within the gene promoter [15, 16]. Only a poly-G length of eight or fewer nucleotides, for example, was permissive for transcription of the genes encoding the *T. pallidum* repeat (Tpr) E, G, and J putative porins [16], while the poly-G associated with *tp0126* gene (encoding an OmpW homolog) and located between the −35 and −10 consensus sequences of the *tp0126* promoter, allowed optimal gene transcription when its length brought the overall distance between the −35 and −10 sites to 17 nucleotides, which is known to be optimal for RNA polymerase binding [15].

Transcriptional changes induced by stochastic expansion and contraction in length of repetitive sequences, such as homomonomeric or homodimeric repeats are collectively known as phase variation, a mechanism used by pathogenic bacteria to rapidly create phenotypic diversity within a population. When this process influences the expression of surface antigens, like in the case of Tp0126 or the Tpr proteins [15, 16], it could facilitate immune evasion or perhaps foster adaptation to diverse host microenvironments. An example is the variable expression of opacity (Opa) proteins in *Neisseria meningitidis*, reported to change the pathogen’s tropism for human epithelium, endothelium, and phagocytic cells [17]. In *T. p. pallidum*, the presence of a poly-G upstream of an annotated ORFs could, therefore, be an indicator that the gene undergoes phase variation, particularly if the poly-G localizes within the experimentally determined or predicted gene promoter. Additionally, like in the case of the Tp0126 ORF, the experimental assessment of the poly-G position in relation to the gene transcriptional start site (TSS) allowed us to redefine the length of the ORF and identify a putative NH_2_-terminal cleavable signal peptide previously embedded within the larger reading frame originally but mistakenly annotated. Such finding supported Tp0126 as a novel OMP, and allowed its identification as an OmpW homolog of the syphilis spirochete. A poly-G is also reported upstream of the *tp1031* gene, encoding the Tpr protein, TprL, which is significantly conserved among syphilis strains and subspecies and hence a possible vaccine candidate.

In the current study, we investigated the role of the *tprL*-associated poly-G in transcription of this gene after redefining the boundaries of this ORF. Additionally, we compared *tprL* transcription in *T. p. pallidum* to an isolate of *T. p. pertenue* (the Gauthier strain), the spirochete agent of the endemic treponematosis yaws [18]. Although nearly identical to the syphilis spirochete at the genomic level, *T. p. pertenue* strains carry a 378-bp deletion that eliminates the poly-G upstream of the *tprL* gene as well as the annotated gene start codon (SC), providing a naturally-occurring mutant for our studies, given that the agents of human treponematoses cannot be genetically altered. Finally, we compared the humoral response to TprL in rabbits and patients infected with the agents of syphilis and yaws to gain insight into whether TprL is produced by the yaws agent.

## Materials and Methods

### Ethics Statement

Only male New Zealand White (NZW) rabbits ranging from 3.5-4.5 Kg were used in our studies. Specific pathogen-free (SPF; *Pasteurella multocida*, and *Treponema paraluiscuniculi*) animals were purchased from Western Oregon Rabbit Company (Philomath, OR) and housed at the University of Washington (UW) Animal Research and Care Facility (ARCF). Care was provided in accordance with the procedures described in the Guide for the Care and Use of Laboratory Animals [19] under protocols approved by the UW Institutional Animal Care and Use Committee (IACUC; Protocol # 4243-01, PI: Lorenzo Giacani). However, because only random animals were tested for *T. paraluiscuniculi* infection by the vendor, all rabbits were bled and tested with a treponemal (FTA-ABS, Trinity Biotech, Bray, Ireland) and a non-treponemal test (VDRL, Becton Dickinson, Franklin Lakes, NJ) upon arrival at the ARCF and prior to use. Both tests were performed according to the manufacturer instructions, with the exception that a secondary FITC-labelled goat anti-rabbit IgG was used instead of the anti-human secondary for the FTA-ABS test. Only seronegative rabbits were used.

Human serum specimens from yaws- and syphilis-infected patients were obtained as de-identified samples and did not require IRB approval. More specifically, twenty-five sera from serologically confirmed (RPR, TPPA and/or EIA) syphilis-infected patients were collected by the King County Public Health (KCPH) laboratory at Harborview Medical Center in 2020. All but one sample were tested by RPR and yielded titers ranging between 1:4 and 1:512. Sera from serologically confirmed (RPR, TPPA and/or EIA) yaws-infected patients (n=25) were collected by Dr. Oriol Mitjá while leading the WHO-sponsored yaws elimination campaign in Lihir Island, Papua New Guinea [20].

### Experimental infections and nucleic acid extraction

Three *T. p. pallidum* strains (Nichols, Chicago, and Seattle 81-4), and one *T. p. pertenue* (Gauthier) strain were propagated by means of intratesticular infection as previously reported [21]. Rabbits were infected with 2 x 10^7^ *T. pallidum* per testis for Nichols and Chicago, and 5 x 10^6^ organisms per testis for Seattle 81-4, and Gauthier. Treponemes were harvested at peak orchitis (approximately day 10 post-infection for the Nichols and Chicago strains; day 20 for Seattle 81-4, and Gauthier) to recover organisms prior to immune clearance. Briefly, testes were minced in 10 ml of saline and shaken for 5 min. Suspensions were spun for 10 min at 1,000 rpm in a 5430 Eppendorf centrifuge (Eppendorf, Hauppauge, NY) to remove host cellular debris. For RNA and DNA isolation, 1-ml aliquots were spun for 30 min at 12,000 rpm at 4°C and the pellets resuspended in 400 μl of Trizol buffer (Thermo Fisher Scientific, Waltham, MA) or 400 μl of DNA lysis buffer (5 mM Tris, pH 8.0; 50 mM EDTA; 0.25% SDS), respectively. For the analysis of the length of the poly-G associated to the *tprL* (*tp1031*) ORF, DNA was isolated as previously described [22] using the QIAamp DNA Mini Kit (Qiagen Inc., Chatsworth, CA). RNA extraction was performed following Trizol manufacturer’s instructions. Prior to reverse transcription, total RNA samples were treated with DNase I (Thermo Fisher Scientific). DNA-free RNA was checked for residual DNA contamination by qualitative amplification using primers specific for the *tp0574* gene encoding the 47 kDa lipoprotein (Sense primer 5’-cgtgtggtatcaactatgg, and antisense primer 5’-tcaaccgtgtactcagtgc, conserved in all strains) as already described [23]. Reverse transcription (RT) of total RNA was performed using the Superscript III First Strand Synthesis Kit (Thermo Fisher Scientific) with random hexamers according to the provided protocol. cDNA samples were diluted 1:5 with molecular grade water and stored in single-use aliquots at −80°C until use.

Intradermal (ID) experimental infections to assess development of humoral immunity to recombinant TprL (TprL) over time were performed on a total of four rabbits. Two rabbits were infected with the Nichols strain in six sites on their shaved backs, and two with the Gautier strain immediately after harvesting from a routine intratesticular strain passage. Each site received 10^6^ spirochetes. Blood was collected from these rabbits at regular intervals for ~90 days. Extracted serum was heat-inactivated at 56°C for 30 min and stored at −20°C until use.

### Quantification of *tprL* message by RT-qPCR

A relative quantification protocol using external standards was used to analyze the *tprL* message level at the time of bacterial harvest from rabbit testes. This approach normalizes the amount of message from the *tprL* gene to that of the *tp0574* gene, used as housekeeping gene. To obtain the standards, sequences of the *tprL* and *tp0574* genes were amplified from Nichols DNA using primers conserved across all strains and cloned into a pCRII-TOPO vector (Thermo Fisher Scientific). For *tprL*, sense and antisense primers 5’-ataagaat<u>gcggccgc</u>ggtggtttcccatttggaagg and 5’-ataagaat<u>gcggccgc</u>caagtagtctgtaagctgcctg (amplicon size: 295 bp) were used, while for *tp0574*, the same primers listed in the above paragraph were used (amplicon size: 313 bp). The *tp0574* amplicon was directly cloned into the vector TA site, while the *tprL* amplicon was cloned using the NotI restriction site (underlined in primer sequences. Resulting construct was linearized by EcoRV digestion, and standard curves were generated by serially diluting the plasmid (tenfold) over the 10^6^-10^0^ copies/μl concentration range. The threshold value for the maximum acceptable error associated with a standard curve was set to 0.05. Amplification reactions and data collection were carried out on a Roche LightCycler (Basel, Switzerland). All reactions were performed following the manufacturer’s instructions with the Roche FastStart Universal SYBR Green Master Kit (Roche). The same primers reported above for *tp0574* and *tprL* (but without restriction tags in the latter case) were used for the qPCR. Amplifications were performed with three microliters of the final cDNA preparation in quadruplicate. Amplification conditions for *tp0574* were: annealing at 60°C for 8 sec following hot start, and extension for 13 s at 72°C. Amplicon melting temperature was 88°C. Amplification conditions for *tprL* were: annealing at 62°C for 6 s following hot start, and extension for 12 s at 72°C. Amplicon melting temperature was 90°C. Differences between levels of *tprL* expression within strains were compared using Students t-test, with significance set at *p*<0.05.

### Identification of the *tprL* transcriptional start site

Rapid Amplification of cDNA Ends (5’-RACE, Thermo Fisher Scientific) was used to determine the *tprL* gene transcriptional start site (TSS) and infer the location of the *tprL* promoter. 5’-RACE was performed on total RNA from *T. p. pallidum* Nichols Seattle and *T. p. pertenue* Gauthier strains following the kit manufacturer’s instructions. For each strain the procedure was carried on in duplicate using the same template RNA. Briefly, for the initial reverse transcription step, 1 μg of sample RNA and 2.5 pmoles of a first *tprL*-specific antisense primer (5’-gtcaggtacgcgttgtagca) were used for reverse transcription, which was followed by dC-tailing of the cDNA. The subsequent amplifications were performed using five microliters of dC-tailed cDNA in 50 μl final volume containing 2.5 units of GoTaq polymerase (Promega), 200 μM of each dNTP, 1.5 mM of MgCl_2_, and 400 nM of a second *tprL*-specific antisense primer (5’-ggagcgttgcttcaaaagac) annealing upstream of the one used for first-strand synthesis and the provided Abridged Anchor Primer. Cycling parameters were initial denaturation (94°C) and final extension (72°C) for 10 min each. Denaturation (94°C), annealing (60°C) and extension (72°C) steps were carried on for 1 min each for a total of 45 cycles. PCR products were purified QIAquick PCR Purification Kit (Qiagen) and cloned into the pCRII-TOPO-TA vector (Thermo Fisher Scientific) according to instructions. For each cloning reaction, plasmid DNA from at least ten colonies was extracted using the Plasmid Mini Kit (Qiagen) and sequenced with vector-specific sense and antisense primers. Sequence data were analyzed using Bioedit, available at https://bioedit.software.informer.com/.

### Analysis of the *tprL*-associated poly-G length

A DNA fragment of 292 bp containing the *tprL*-associated poly-G repeat was amplified for fluorescent fragment length analysis (FFLA), a method already used to evaluate the variability of poly-G tracts upstream of *T. p. pallidum* genes among and within isolates [15]. Briefly, amplification was performed using a 6-NED-labeled sense primer (5’-cacggggcgatacaaaactc) and an unlabeled antisense primer (5’-gtttcttccctcccgacccatttcatt). Amplifications were performed in 50 μl final volume using 2 U of AccuPrime *Pfx* Polymerase (Thermo Fisher Scientific) and 100 ng of DNA template in each reaction. Mix was also supplied with primers, MgSO_4_, and dNTPs at final concentrations of 300 nM each, 1 mM, and 300 μM, respectively. Amplifications were carried on for 45 cycles, with denaturation (94°C), annealing (60°C) and extension (68°C) times of 30 sec, 30 sec, and 1 min, respectively. Initial denaturation (94°C) and final extension (68°C) steps were of 10 min each. For each strain, two independent amplifications were performed using the same template DNA. Amplification products were purified using the QIAquick PCR Purification Kit (Qiagen). Concentrations were measured spectrophotometrically and all samples diluted to 0.2 ng/μl final concentration. One microliter of each sample was mixed with 15.4 μl of highly deionized formamide and 0.1 μl of HD400 ROX-labeled DNA size marker (both reagents from Thermo Fisher Scientific).

Samples were transferred to a 96-well plate and denatured by incubation at 94°C for 2 min, and loaded onto an ABI3730xl DNA analyzer (Thermo Fisher Scientific) to be separated by capillary electrophoresis. Electropherograms were analyzed using the GeneMapper 4.0. Data on amplicon length (determined by comparison to the ROX-labeled marker) and intensity (measured as area under a peak) were collected to evaluate the proportion of amplicons with poly-G’s of different length within each sample. For each amplification, FFLA was performed in triplicate. For data analysis, the sum of the area underneath all peaks generated by amplicons with the same number of G’s was divided by the total area underneath all peaks.

### GFP reporter assay

The *tprL* promoter was amplified using the sense and antisense primers 5’-ccccctgtctacctgagga and 5’ - gcatggtgcagttccttccc, respectively and cloned into the pGlow-TOPO TA vector (Thermo Fisher Scientific), carrying a promoter-less GFP reporter gene. Primers were designed to include the putative TprL start codon as the first codon of the GFP ORF. Amplification was performed in 50 μl final volume using 2 U of GoTaq Polymerase (Promega) and 100 ng of DNA template in each reaction, and carried out for 45 cycles, with denaturation (94°C), annealing (60°C) and extension (68°C) times of 30 sec. Initial denaturation (94°C) and final extension (68°C) steps were of 10 min each. Amplification products were cloned directly into the pGlow-TOPO vector according to the manufacturer’s instructions. Amplicon included 90 nt upstream of the poly-G tract to include the *tprL* promoter, and a putative ribosomal binding site (RBS, GGAG) located 4 nucleotides upstream of the TprL predicted start codon. With the exception of the start codon, no other TprL codons were present in the constructs. Expression of GFP from these constructs resulted in the addition of nine extra amino acids to the actual GFP peptide, encoded by the TprL start codon and eight additional vector-encoded. In total, two different constructs were obtained for the *tprL* promoter, with poly-G repeats 8 and 10 nt long. A construct containing the *lac* promoter upstream of the GFP gene was used as a positive control. As a negative control, to determine background fluorescence, the *tp0547* ORF fragment (the same used for message quantification purposes, see above) and not predicted to harbor a promoter or a ribosomal binding site was inserted upstream of the GFP coding sequence of the pGlow-TOPO vector. All constructs were sequenced on both strands to verify sequence accuracy and correct insert orientation using sanger seqeuncing. Constructs were then used to transform TOP-10PE *E. coli* cells (Thermo Fisher Scientific) which do not carry the *lacI* repressor gene. For GFP fluorescence measurements, cells transformed with the various constructs were inoculated from a plate into 4 ml of LB-ampicillin (100 μg/ml) broth and grown at 37°C for 4 hr. Optical density (OD_600_) of all cultures was then measured using a biophotometer (Eppendorf) and cultures were diluted to identical optical density (0.5 Absorbance Units, AU). Subsequently, OD_600_ and fluorescence were recorded in parallel until cultures reached an OD_600_ of ~2 AU. For fluorescence readings, 400 μl of culture were centrifuged for 4 min at full speed on a tabletop centrifuge and resuspended in an equal volume of phosphate buffered saline (PBS). Cells were then divided in three wells (100 μl/well) of a black OptiPlate-96F (Perkin Elmer, Boston, MA) for top fluorescence reading. Excitation and emission wavelength were 405 and 505 nm, respectively, and readings were performed in a BioTek Synergy Microplate Reader (BioTek, Winooski, VT). Reported data represent fluorescence (expressed in Arbitrary Units, A.U.) normalized to the optical density of the culture. Background fluorescence values were obtained using *E. coli* cells transformed with the reporter vector containing the promoterless *tp0574* ORF fragment. Differences in levels of fluorescence between cultures were compared using Student’s t-test, with significance set at *p*<0.05.

### ELISA with recombinant TprL and Tp0574 antigens

A *tprL* gene, devoid of signal peptide and codon-optimized for expression in *E. coli* was synthesized by GenScript. The gene was then subcloned into the pET28a(+) vector, between the BamHI and XhoI sites. Recombinant TprL spanned 489 amino acids. The *tp0574* gene was amplified using the primers sense 5’-tgtggctcgtctcatcatga and antisense 5’-ctgggccactaccttcgcac. Amplification was performed in 50 μl final volume using 2 U of GoTaq Polymerase (Promega) and 100 ng of Nichols DNA template. Mix was also supplied with primers, MgSO_4_, and dNTPs at final concentrations of 300 nM each, 1.5 mM, and 300 μM, respectively. Amplifications were carried on for 45 cycles, with denaturation (94°C), annealing (60°C) and extension (68°C) times of 30 sec, 30 sec, and 1 min, respectively. Initial denaturation (94°C) and final extension (68°C) steps were 10 min each. Amplicon was cloned directly the pEXP-5-NT/TOPO vector (Thermo Fisher Scientific). Constructs were sequenced prior to expression to ensure lack of amplification errors in the transgene as well as correct orientation into the vector. For protein expression, transformed *E. coli* Rosetta2 DE3 pLysS BL21 derivative cells (Sigma-Aldrich, ST. Louis, MO) were grown at room temperature in auto-inducing media according to Studier *et al.* [24] and harvested after 3 days of incubation. Purification was performed by nickel affinity chromatography under denaturing conditions using the Ni-NTA Agarose gravity chromatography System (Qiagen). Inclusion bodies containing insoluble recombinant proteins were isolated by successive rounds of sonication and centrifugation, then resuspended in 1X binding buffer (0.5 M NaCl, 20 mM Tris-HCl, 5 mM imidazole, pH 7.9) containing 6 M urea. After 1 h incubation in ice, suspensions were centrifuged, and supernatants passed through a 0.45 μm filter. Purification was followed by dialysis against PBS. Products were tested for size and purity by SDS-PAGE and quantified using a Bicinchoninic Acid Assay kit (Pierce, Rockford, IL).

Purified recombinant TprL in PBS and recombinant Tp0574 (the 47 kDa lipoprotein, as a positive control antigen), were used to coat the wells of a 96-well flat bottom EIA/RIA microplates (Corning LifeSciences, Corning, NY). Plates, containing 15 picomoles/well of TprL or Tp0574 protein in 50 μl were incubated at 37°C for 2 h and subsequently at 4°C overnight to induce antigen binding to the test wells. Wells were then washed three times with PBS containing 0.05% Tween-20 (Sigma-Aldrich), blocked by incubation overnight at 4°C with 200 μl of 3% nonfat milk-PBS/well and washed again the next morning. Ten microliters of each serum (either from *T.p. pallidum* or *T.p. pertenue* infected animals or patients) were diluted 1:20 in 1% nonfat milk-PBS and 100 μl dispensed into wells. Sera were incubated over night at room temperature. Wells were then washed three times with PBS containing 0.05% Tween-20 (Sigma-Aldrich). One hundred microliters of secondary antibody (alkaline phosphatase-conjugated goat anti-rabbit IgG or goat anti-human IgG, both from Sigma-Aldrich) diluted 1:2,000 in 1% nonfat milk-PBS were then added to each well and the plates incubated for additional 3 h at room temperature before repeating the washing step. After addition of 50 μl of 1 mg/ml para-nitrophenyl phosphate (Sigma-Aldrich) to each well, plates were developed for 45 min, and read at 405 nm on a BioTek Microplate reader. The mean of background readings (from no antigen control wells) was subtracted from the mean of triplicate experimental wells for each serum.

### Genome-wide analysis of the poly-G sequence variability in *T. p. pallidum*

A previously performed analysis revealed several poly-G tracts (≥ 8 nt) [15] distributed throughout the *T. p. pallidum* Nichols strain genome [25] associated to as many genes. To investigate how many of those elements are variable in the *T. p. pallidum* strains used here, and hence possibly affecting gene expression at the transcriptional or translational level, we applied the same FFLA technique described above for the *tprL* poly-G to each of these homopolymeric tracts. Primers are reported in Table 1. Forward primers were labelled with different fluorophores (FAM, HEX, or NED) to multiplex three targets at the time. Amplification and separation by capillary electrophoresis were performed as described above for the *tp1031* gene.

**Table 1.**
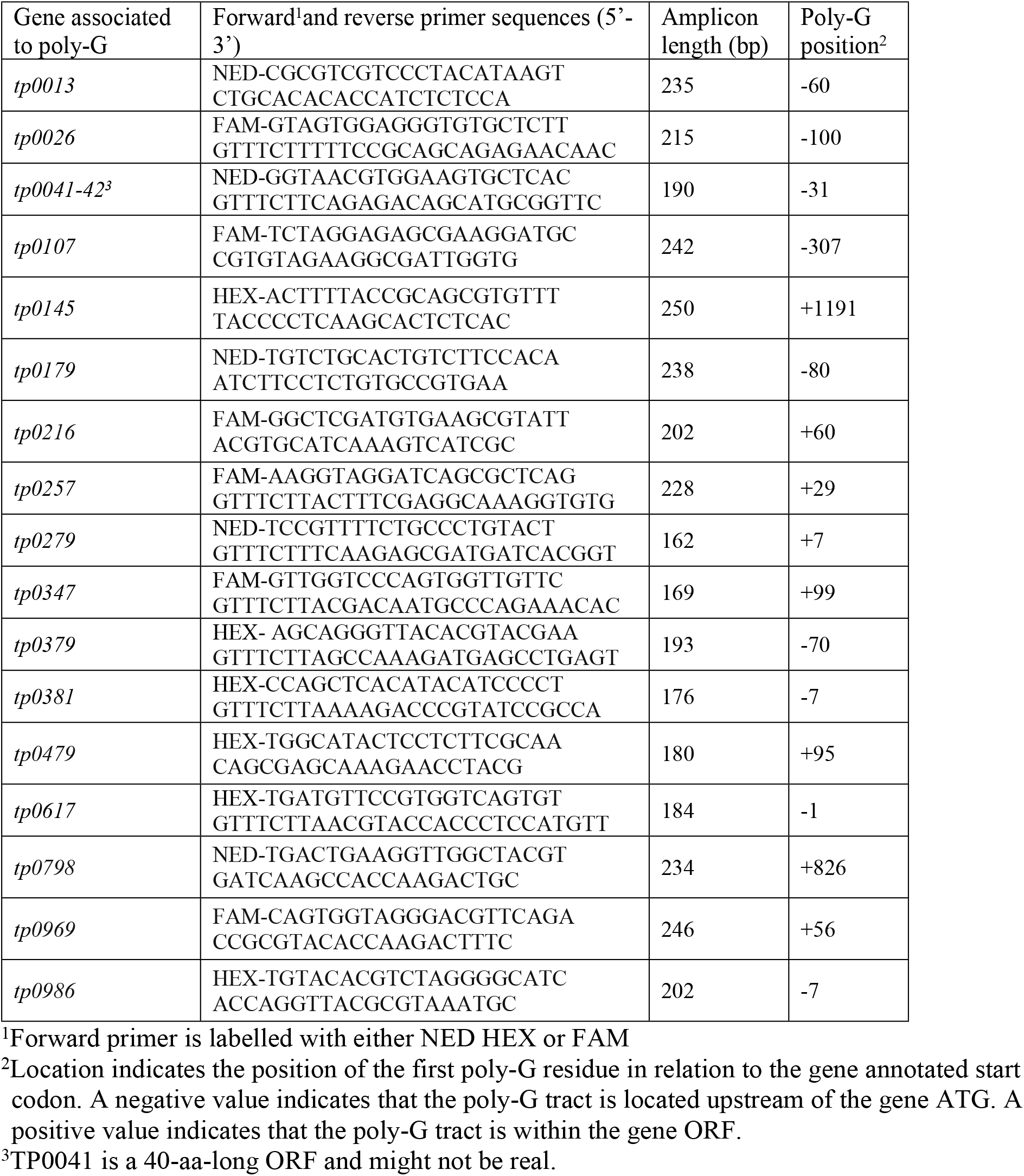
Primers used in this study

### Analysis of the poly-G sequence variability in *T. p. pallidum* and *in silico* analysis of the extended TprL protein

Of the 17 targets analyzed, 88% was found to be variable in length, while only two did not appear to be (Fig.5). *In silico* analysis on the extended TprL ORF to support it as a putative *T. pallidum* OMP was performed using a series of computational tools. Presence of a cleavable signal peptide was predicted by SignalP 4.1 (http://www.cbs.dtu.dk/ervices/SignalP/) [26], PrediSi (http://www.predisi.de/) [27], and LipoP (http://www.cbs.dtu.dk/services/LipoP/) [28]. For OM location, we used CELLO [29], PSORTb 3.0 [30], BOMP [31], HHPRED (https://toolkit.tuebingen.mpg.de/tools/hhpred) [32] and PRED-TMBB (http://bioinformatics.biol.uoa.gr/PRED-TMBB/) [33]. Sequence and structural homology of the TprL ORF to other bacterial proteins was investigated using Phyre2 (http://www.sbg.bio.ic.ac.uk/~phyre2/html/page.cgi?id=index) [34], I-TASSER (https://zhanglab.ccmb.med.umich.edu/I-TASSER/) [35] and LOMETS (https://zhanglab.ccmb.med.umich.edu/LOMETS/) [36] using default parameters.

## Results

### Analysis of *tprL* transcription levels during experimental infection

To examine whether differential expression of the *tprL* gene occurs in *T. pallidum* strains and subspecies, a real-time qPCR assay was developed to quantitate *tprL* message in treponemal strains harvested at the same time during experimental infection (peak orchitis). That approach normalizes the *tprL* message level to that of the *tp0574* gene as previously described [37]. Quantification data showed that *tprL* mRNA is variably expressed in the isolates analyzed here (Fig.1). Compared to *T. p.* subsp. *pertenue* (Gauthier strain), all *T. p.* subsp. *pallidum* strains (Nichols, Chicago, and Seattle 81-4) showed a significantly higher level of *tprL* mRNA (p<0.05; Fig.1). *tprL* mRNA levels detected in Nichols and Chicago were not significantly different, while the Seattle 81-4 strain showed the higher message level of this gene (Fig.1) among the syphilis isolates. This result supported the existence of mechanisms affecting *tprL* transcription.

**Figure 1.**
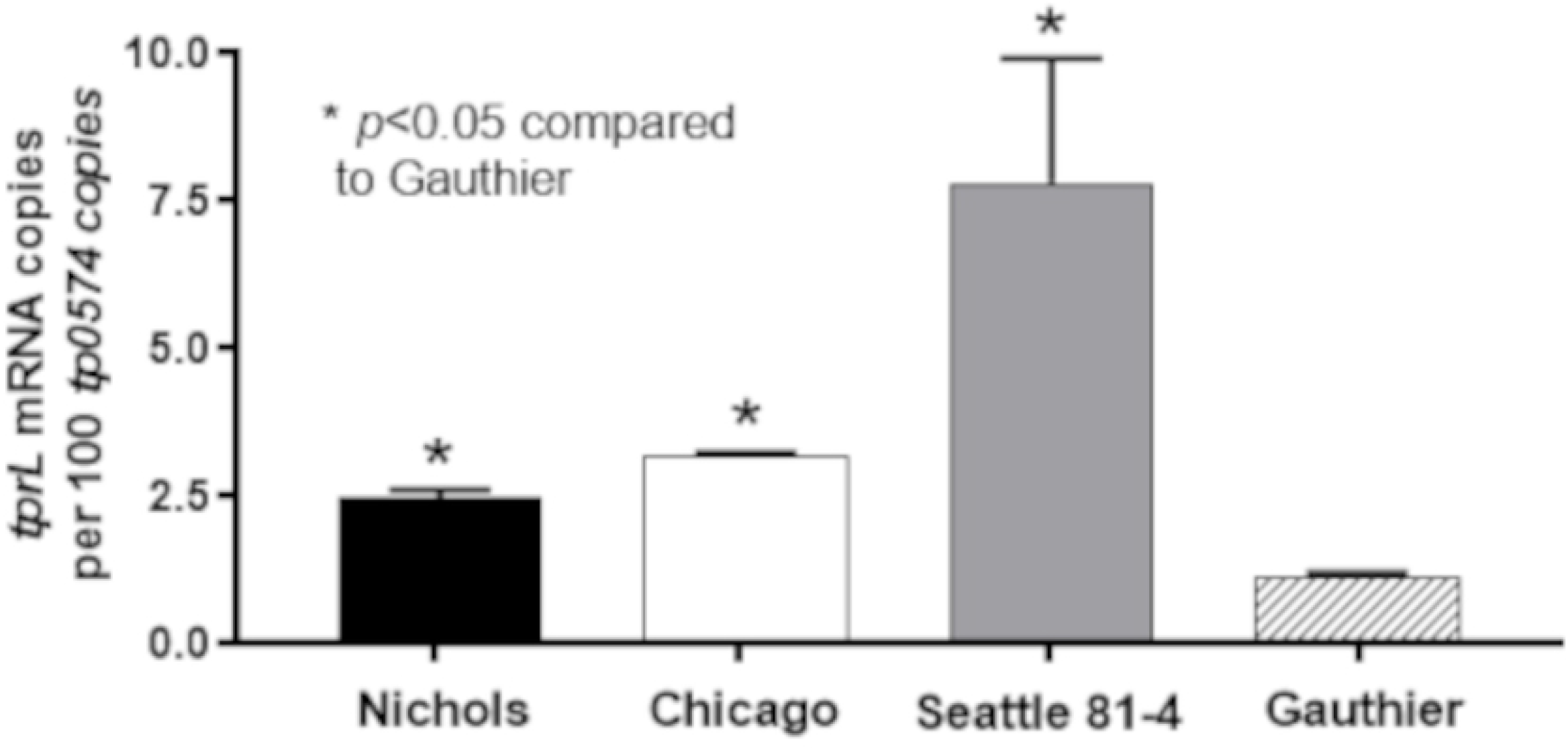
*tprL* mRNA levels normalized to the *tp0574* message in the *T. p. pallidum* Nichols, Chicago and Seattle 81-4 strains and in the in the *T. p. pertenue* Gauthier strain harvested at peak orchitis post-IT inoculation. Asterisk (*) indicates a significant difference (*p*<0.05) compared to the Gauthier strain.

### Identification of the *tprL* transcriptional start site and analysis of variability of the *tprL*-associated poly-G

Upon performing 5’-RACE with *T. p. pallidum* RNA, the *tprL* TSS was identified to be the fourth nucleotide downstream of the poly-G (Fig.2A). Such finding was similar to what previously reported for other *tpr* genes paralogous of *tprL* that also carry a poly-G of varying length upstream of their TSS [16]. We then re-assessed the TprL protein annotation. Analysis of the sequence downstream of the newly identified TSS allowed us to predict a putative ribosomal binding site (RBS) and an alternative start codon (SC) for TprL (Fig.2A). Such prediction extended the previously annotated TprL ORF by 88 codons. More importantly, this additional sequence was predicted to contain a cleavable signal peptide (aa 1-25, underlined in Fig.2A) by PrediSi, LipoP, and SignalP, which is necessary for OMP sorting to the bacterial surface [38, 39]. 5’-RACE was also performed using Gauthier total RNA but did not yield any reproducible result, suggesting that the TSS of the transcript carrying the *tprL* message in Gauthier was not in proximity of the primers used. Given the location of the poly-G in the *tprL* promoter region, we then investigated whether this poly-G showed length variability *in vivo* within each syphilis strain studied here. To this end, we used a FFLA method based on the amplification of the poly-G repeat with a fluorescent primer and subsequent size separation on a genetic analyzer. The results are shown in Fig.2B. FFLA showed that the length of the *tprL*-associated poly-G varies *in vivo* within each isolate. More specifically, the *tprL*-associated poly-G was shown to contain homopolymeric tracts varying from 7 to 11 Gs, even though the vast majority of the fragments contained 9 Gs. Poly-G length distribution between Chicago and Seattle 81-4 strains was found to be more similar, with a comparable percentage of amplicons containing 8, 9, 10, and 11 Gs, respectively, while no amplicons containing 7 Gs were detected in either strain (Fig.2B).

**Figure 2.**
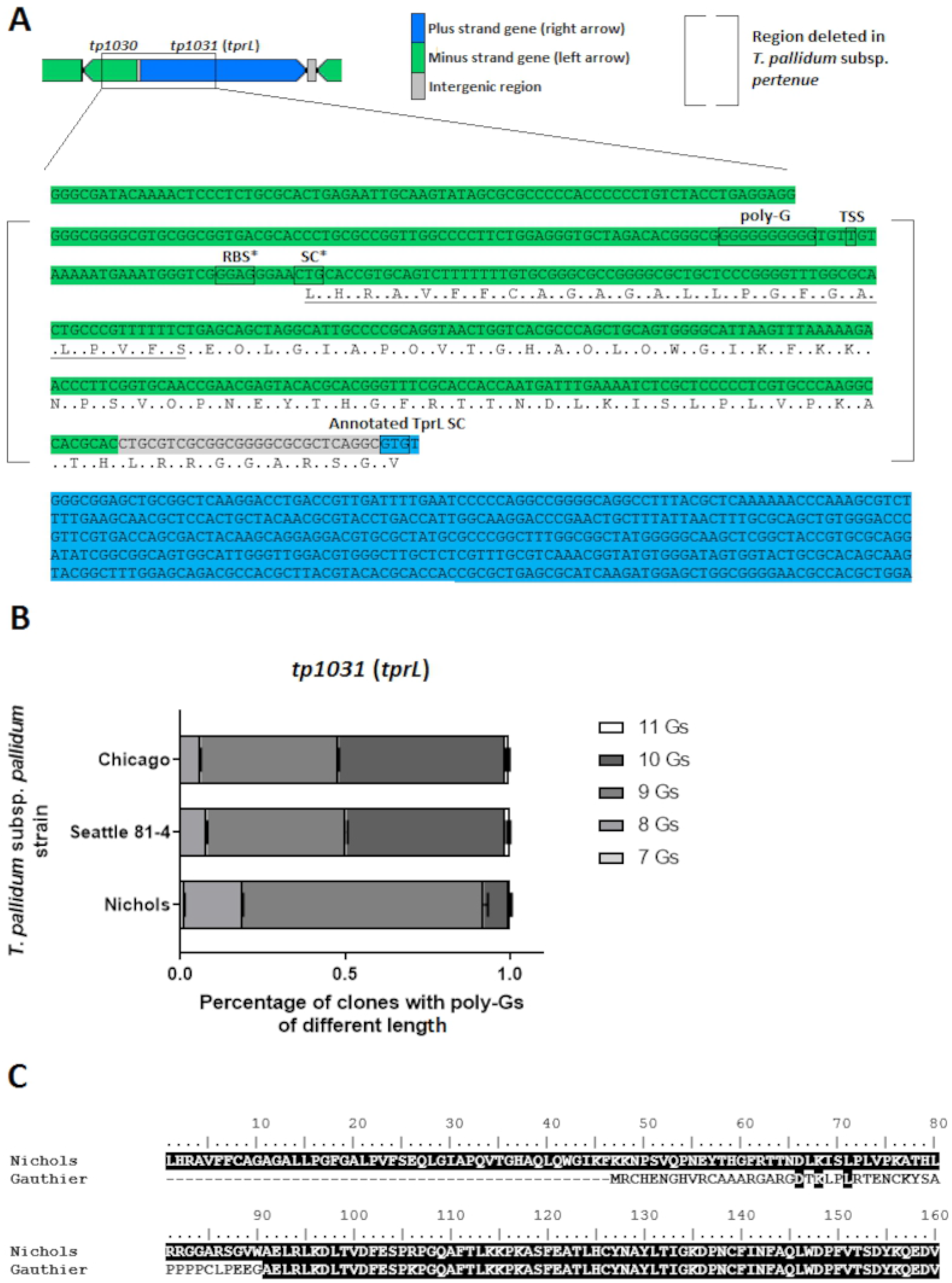
(**A**). Cartoon depicting the *tprL* ORF as originally annotated in the *T. pallidum* Nichols strain [25] (blue) showing the location of the homopolymeric G tract (poly-G), the experimentally determined TSS. Asterisks indicate the newly predicted (*) TprL ribosomal binding site (RBS) and start codon (SC) based on the TSS identification. According to this model, the TprL ORF encompasses 88 additional amino acid residues located upstream of the annotated SC. Within these residues, a cleavable signal peptide (underlined) is predicted by three independent programs (SignalP, PrediSi and LipoP). (**B**). Distribution of poly-G lengths in the Nichols, Chicago, and Seattle 81-4 treponemes at the moment of bacterial harvest determined by FFLA. (**C**) Comparison of the TprL NH_2_-termini of *T. p. pallidum* and *T. p. pertenue* (Nichols and Gauthier strains, respectively).

### Role of poly-G length in transcription

To investigate whether poly-G repeats of different length would affect the activity of the *tprL* promoter, we adopted an *E. coli*-based heterologous system that allows monitoring of expression of a vector-encoded GFP reporter gene placed under the control of the *tprL* promoter with poly-Gs of different length. This approach was previously used to evaluate the role of poly-G repeats in transcription of the *tprF, I, E, J*, and *tp0126* genes, which also encode *T. p. pallidum* putative OMPs [15, 16]. For this study, two different *tprL* promoters (with poly-Gs of 8 and 10 nt, respectively) were tested along with positive and negative controls (the *lac* promoter, and a promoter-less reporter vector, respectively). Results (Fig.3) showed that higher GFP fluorescence signal was detected when the *tprL* promoter carried a poly-G of eight residues compared to 10 G residues, which induced a fluorescence signal slightly above background but not significantly different. This result supports the hypothesis that *tprL* expression is influenced by phase variation.

**Figure 3.**
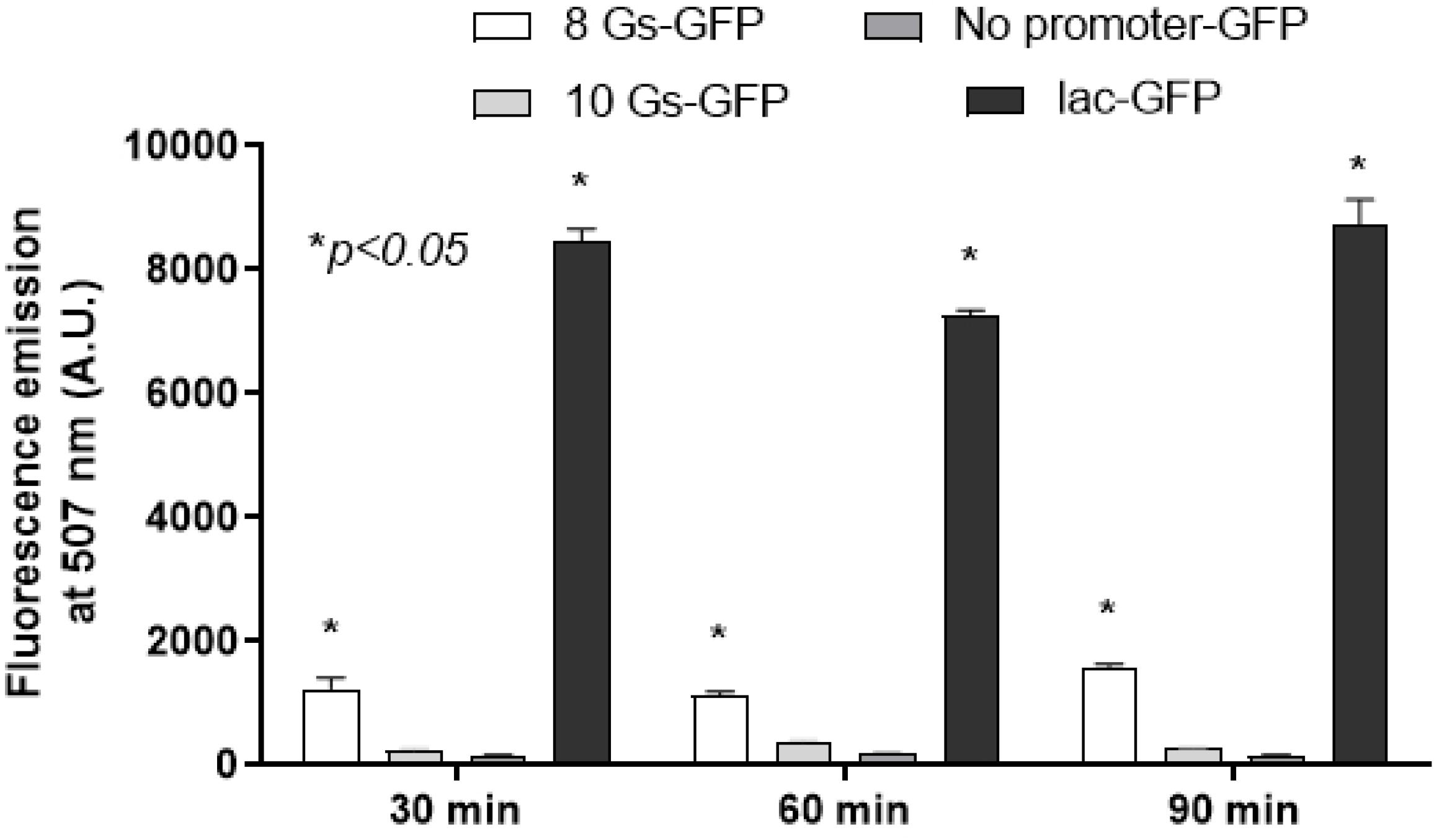
Analysis of the effect of poly-G length on *tprL* transcription. Graph shows fluorescence induced in *E. coli* TOP-10PE cells transformed with a pGLow-TOPO vector where GFP transcription is under control of the tprL promoters with poly-G tracts of different length (8 and 10 nt). A *lac* promoter –GFP construct was used as a positive control. The *lac* promoter is recognized by σ^70^ and the *E. coli* strain used for this assay does not carry the gene that encodes the LacI repressor. Background fluorescence collected from *E. coli* cells transformed with a pGLow-TOPO vector that carries a fragment of *T. pallidum Tp0574* ORF with no promoter is also shown. Asterisk (*) indicates significance compared to background fluorescence level (No promoter-GFP sample).

### Humoral response to TprL during experimental and natural infection

Given the apparent lack of elements able to drive translation of TprL in yaws strains, we hypothesized that only syphilis-infected rabbits and patients would develop humoral immunity to TprL when compared to their yaws-infected counterparts. Therefore, we evaluated sera from longitudinally infected rabbits and syphilis patients by ELISA using recombinant TprL. ELISA results using animals infected with the Nichols strain showed limited but detectable reactivity to TprL, which developed relatively late (day 30) post-inoculation (Fig.4A). A more pronounced response to TprL was seen in naturally infected patient samples (Fig.4B). As expected, the vast majority of cases showed a significantly higher reactivity to the Tp0574 antigen, used as a positive control (Fig.4A-B). Conversely, no reactivity to TprL was seen in sera from yaws-infected animals or patients, while reactivity to Tp0574 remained readily detectable (Fig.4C-D). These results suggest that the deletion naturally occurring in yaws isolates upstream of the TprL ORF might abolish protein expression in this subspecies.

**Figure 4.**
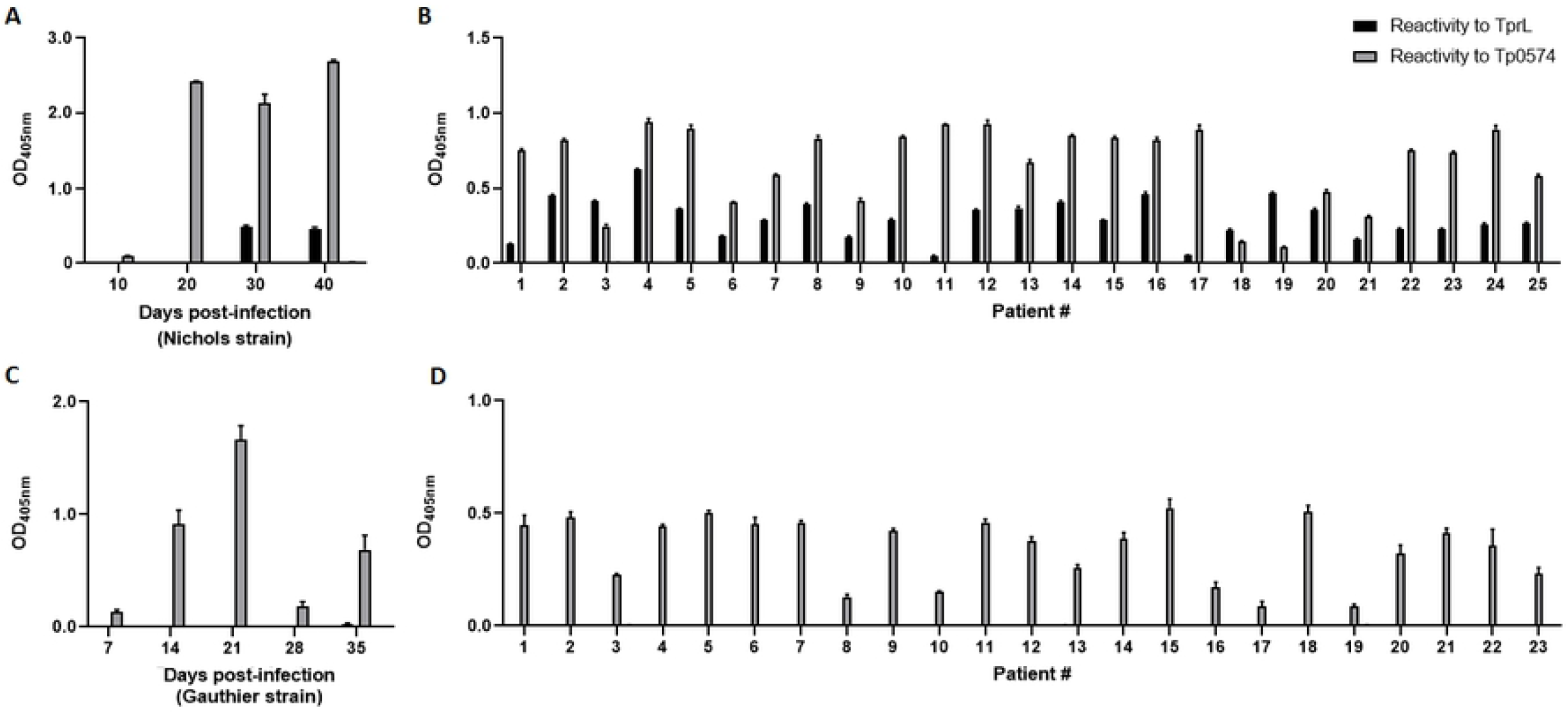
Humoral reactivity to TprL in experimental and clinical samples. (**A**) Pooled sera from rabbits (n=2) infected with *T. p. pallidum* Nichols strain, and (**B**) individual sera from syphilis-infected patients. (**C**) Pooled sera from rabbits (n=2) infected with *T. p. pertenue* Gauthier strain, and (**D**) individual sera from yaws-infected patients. Optical density from test sera in absence of target antigen was used for background subtraction.

## Discussion

The high global prevalence of syphilis and its resurgence in high-income nations argues in favor of deepening our knowledge of syphilis pathogenesis with the goal of better understanding *T. p. pallidum* virulence mechanisms, particularly with regard to its ability to persist in the host in absence of treatment [9]. Such knowledge might help devise better strategies for disease control and accelerate vaccine development. Previous studies have clearly shown that antigenic variation of the OMP TprK plays a major role in *T. p. pallidum* virulence and survival *in vivo* [40–42]. Phase variation is a second, distinct mechanism that has more recently been implicated in *T. pallidum* virulence based upon our studies. Phase variation allows rapid and reversible ON/OFF switching of gene expression at the transcriptional or translational level to generate phenotypic antigenic diversity during infection. This mechanism is mediated by rapid changes in length of DNA repeats such as homopolymeric tracts due to slipped-strand mispairing during replication. Such elements affect transcription or translation when located within a gene promoter or an ORF, respectively. In pathogens such as *N. meningitidis* or *H. pylori*, for example, phase variation influences expression of determinants involved in immune evasion as well as in adaptation to different host microenvironments [43–45].

Here, we studied the poly-G associated to the *tprL* gene to assess whether this gene could also undergo phase variation. Given that TprL is also a putative OMP and significantly conserved among strains and subspecies of *T. pallidum*, this study is relevant to inform ongoing vaccine development efforts. By identifying the TSS of this gene, we first confirmed that the *tprL* transcript begins over two hundred nucleotides upstream of the annotated protein start codon [25]. This finding supports that the current annotation of the TprL protein should be revised, as the TprL coding sequence likely includes 88 additional NH2-terminal amino acids, encoded by nucleotides currently annotated as part of the *tp1030* gene (Fig.2A). Interestingly, this additional TprL sequence is strongly predicted to contain a cleavable signal peptide, which further supports TprL as a putative OMP. Previous attempts to identify a signal peptide on the shorter TprL protein were not successful [38], even though the same studies overall supported TprL as an OMP based on *in silico* structural homology analysis. Overall, our findings support the necessity to revise the annotation of *T. p. pallidum* genome using high throughput RNA-seq approaches, which will provide a more precise annotation of the protein-encoding genes, intergenic regions, and organization of ORFs in operons. Necessity to re-annotate based on experimental data was also highlighted by our work on the *tp0126* gene, whose signal peptide could be predicted only after determination of the gene TSS exactly as for *tprL* [15].

In the case of TprL, the newly predicted start would be a CTG codon, based on the location of the RBS. Although CTG is not among the most commonly utilized start codons in bacteria, this is not an unusual finding in spirochetes. Bulach *et al.* [46], in fact, reported that in *Leptospira* serovars the frequency of CTG use as a start codon ranges between 17-19%. Another finding worth noting is that in spite of a large deletion affecting its upstream region (Fig.2A), a low transcription level for *tprL* could be detected in the *T. p. pertenue* Gauthier strain (Fig.1). It is unclear where the genetic elements responsible for generating this transcript reside in the Gauthier genome, as our attempts to identify them through 5’-RACE failed. However, more importantly, because no humoral reactivity to TprL was seen in yaws-infected rabbits or patients, it is possible that the TprL message is not translated in Gauthier and more generally, the yaws subspecies as a whole. Alternatively, if protein synthesis does occur in yaws treponemes, it might not generate enough antigen to induce a detectable humoral response. Such a finding would have direct implications for development of diagnostic tools for yaws. Due to the aforementioned deletion (shown in Fig.2A), the sequence of the NH_2_-terminal region of TprL is predicted to diverge from that of syphilis isolates (Fig.2C). Such difference was targeted in the past to try and devise a serological test to differentiate syphilis from yaws infection whenever biological specimens were not suitable for molecular analysis of this region. The possibility of differential diagnosis using TprL-based serological approach, however, could be still feasible based on our results as this antigen might not be synthesized at all during infection with yaws strains. A rapid point-of care test where shared antigens are combined with TprL could indeed help differentiate between these two infections. We acknowledge that the number of patient specimens used in our study is limited and that we purposely included specimens from patients with confirmed infection with *T. p. pertenue* based on molecular analysis, and that our findings will need further experimental confirmation using a much larger cohort of patient samples.

In spite of their many commonalities, the pathogenesis of syphilis and yaws also show remarkable differences [18]. If the yaws spirochetes lack a putative OMP and virulence factor such as TprL, as our results here suggest, further studies should try and address the function of this protein and its possible role in the pathogenesis of these infections. Although we hoped to gain clues on TprL function by conducting structural homology analyses, obtaining a consistent model for this protein remains an elusive task, as prediction softwares (Neff-MUSTER, SparksX, HHpred, and HHsearch, all form the LOMETS package) that identify structural homologs to TprL with a beta-barrel structure do not agree on any particular structural homolog. Predictions include electron transport proteins homologous to the Mtr complex of *Shewanella baltica* [47]; a Type 9 protein translocon homologous to the SprA protein of *Flavobacterium johnsoniae* [48] which however has a molecular mass three times that of TprL, OmpW of *E. coli* [49], which however is significantly smaller in size, and a green fluorescent protein of the hydromedusa *Aqueora victoria.* Further experimental work focusing on the analysis of this protein will shed light on its structure, and provide additional clues to its function and role in disease pathogenesis. Our analysis of the variability of most poly-G tracts found in the Nichols strain genome (Fig.5), strongly suggest that phase variation might be a very strong component of the strategy these spirochetes use to create antigenically distinct cells at the phenotypic level. In our analysis, only one of the poly-G analyzed here did not vary upon performing amplification and separation, but most of the others showed a rather significant variability in terms of length. These data will hopefully provide opportunity to address the role of the genes to which the poly-G is associated in disease pathogenesis and to the biology of this difficult organism.

**Figure 5.**
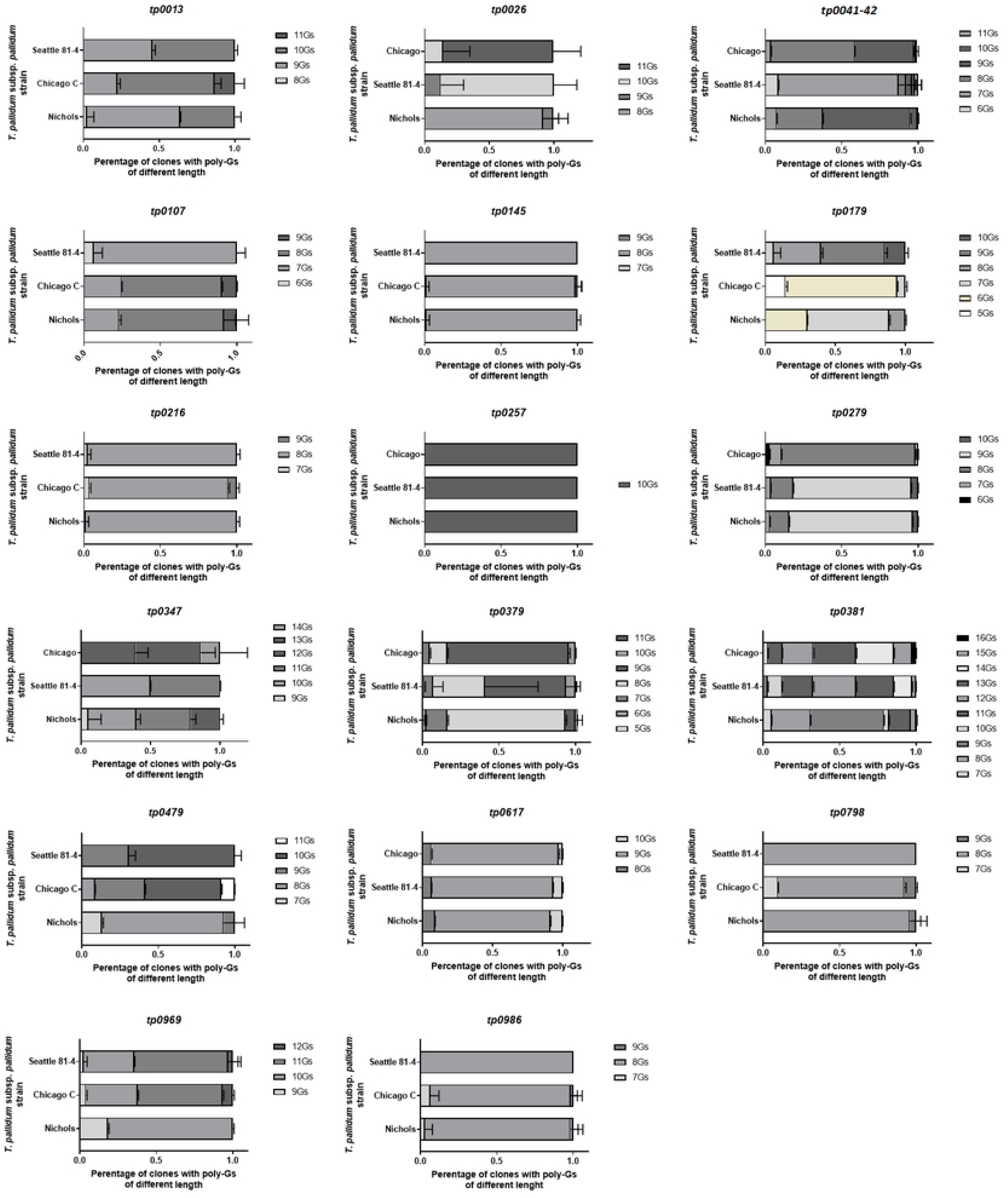
Analysis of poly-G tracts not already studied found throughout the *T. p. pallidum* genome. Because human treponematoses spirochetes have GC-rich genomes (~52.8%), and poly-G tracts are common, we only selected those showing an initial length ≥8 nt.

## Conclusions

Our results support modulation of *tprL* transcription by phase variation and that this gene might not be functional in yaws treponemes. This information might help vaccine design efforts to control syphilis spread.

## Acknowledgments

This work was supported by the National Institute for Allergy and Infectious Diseases of the National Institutes of Health grant numbers R01AI139265 (To J.D. K.) and U19AI144133 Project 2 (Project leader: L.G.; PI: Anna Wald, MD, University of Washington). We are also grateful to Matthew Golden, MD, MPH for help procuring the syphilis patient sera tested in this study. The content of this study is solely the responsibility of the authors and does not necessarily represent the official views of the National Institutes of Health.

## References

1. WHO. Prevalence and incidence of selected sexually transmitted infections *Chlamydia trachomatis*, Neisseria gonorrhoeae, syphilis and Trichomonas vaginalis: methods and results used by WHO to generate 2005 estimates. World Health Organization, Geneva. 2011.

2. Gerbase AC, Rowley JT, Mertens TE. Global epidemiology of sexually transmitted diseases. The Lancet. 1998;351.

3. CDC. 2018 Sexually Transmitted Disease Surveillance. Atlanta, GA: US Department of Health and Human Services: Centers for Disease Control and Prevention. 2019.

4. Savage EJ, Marsh K, Duffell S, Ison CA, Zaman A, Hughes G. Rapid increase in gonorrhoea and syphilis diagnoses in England in 2011. Euro Surveill. 2012;17(29). PubMed PMID: 22835469.

5. Savage EJ, Hughes G, Ison C, Lowndes CM. Syphilis and gonorrhoea in men who have sex with men: a European overview. Euro Surveill. 2009;14(47). PubMed PMID: 19941803.

6. Simms I, Fenton KA, Ashton M, Turner KM, Crawley-Boevey EE, Gorton R, et al. The re-emergence of syphilis in the United Kingdom: the new epidemic phases. Sex Transm Dis. 2005;32(4):220–6. PubMed PMID: 15788919.

7. Tucker JD, Cohen MS. China’s syphilis epidemic: epidemiology, proximate determinants of spread, and control responses. Curr Opin Infect Dis. 2011;24(1):50–5. PubMed PMID: 21150594.

8. Jin F, Prestage GP, Kippax SC, Pell CM, Donovan BJ, Kaldor JM, et al. Epidemic syphilis among homosexually active men in Sydney. Med J Aust. 2005;183(4):179–83. PubMed PMID: 16097913.

9. LaFond RE, Lukehart SA. Biological basis for syphilis. Clin Microbiol Rev. 2006;19(1):29–49. PubMed PMID: 16418521.

10. Goldenberg RL, Thompson C. The infectious origins of stillbirth. Am J Obstet Gynecol. 2003;189(3):861–73. PubMed PMID: 14526331.

11. Nusbaum MR, Wallace RR, Slatt LM, Kondrad EC. Sexually transmitted infections and increased risk of co-infection with human immunodeficiency virus. J Am Osteopath Assoc. 2004;104(12):527–35. Epub 2005/01/18. PubMed PMID: 15653780.

12. Cameron CE, Lukehart SA. Current status of syphilis vaccine development: need, challenges, prospects. Vaccine. 2014;32(14):1602–9. PubMed PMID: 24135571.

13. Bagnoli F, Baudner B, Mishra RP, Bartolini E, Fiaschi L, Mariotti P, et al. Designing the next generation of vaccines for global public health. Omics. 2011;15(9):545–66. PubMed PMID: 21682594.

14. Smajs D, McKevitt M, Howell JK, Norris SJ, Cai WW, Palzkill T, et al. Transcriptome of *Treponema pallidum:* gene expression profile during experimental rabbit infection. J Bacteriol. 2005;187(5):1866–74. PubMed PMID:1063989

15. Giacani L, Brandt SL, Ke W, Reid TB, Molini BJ, Iverson-Cabral S, et al. Transcription of TP0126, Treponema pallidum putative OmpW homolog, is regulated by the length of a homopolymeric guanosine repeat. Infect Immun. 2015;83(6):2275–89. PubMed PMID: 25802057.

16. Giacani L, Lukehart S, Centurion-Lara A. Length of guanosine homopolymeric repeats modulates promoter activity of Subfamily II *tpr* genes of *Treponema pallidum* ssp. *pallidum*. FEMS Immunol Med Microbiol. 2007;51:289–301. PubMed PMID: 3006228

17. Gray-Owen SD. Neisserial Opa proteins: impact on colonization, dissemination and immunity. Scand J Infect Dis. 2003;35(9):614–8. PubMed PMID: 14620144.

18. Giacani L, Lukehart SA. The endemic treponematoses. Clin Microbiol Rev. 2014;27(1):89–115. PubMed PMID: 24396138.

19. Janier M. Ceftriaxone is effective for treating patients with primary syphilis. Sex Transm Dis. 1988;15(1):70. Epub 1988/01/01. PubMed PMID: 3358241.

20. Mitjà O, Houinei W, Moses P, Kapa A, Paru R, Hays R, et al. Mass treatment with single-dose azithromycin for yaws. N Engl J Med. 2015;372(8):703–10. Epub 2015/02/19. doi: 10.1056/NEJMoa1408586. PubMed PMID: 25693010.

21. Baker-Zander SA, Fohn MJ, Lukehart SA. Development of cellular immunity to individual soluble antigens of *Treponema pallidum* during experimental syphilis. J Immunol. 1988;141(12):4363–9. PubMed PMID: 2461990

22. Centurion-Lara A, Arroll T, Castillo R, Shaffer JM, Castro C, Van Voorhis WC, et al. Conservation of the 15-kilodalton lipoprotein among *Treponema pallidum* subspecies and strains and other pathogenic treponemes: genetic and antigenic analyses. Infect Immun. 1997;65(4):1440–4. PubMed PMID: 175151

23. Giacani L, Hevner K, Centurion-Lara A. Gene organization and transcriptional analysis of the *tprJ, tprI, tprG* and *tprF* loci in the Nichols and Sea 81-4 *Treponema pallidum* isolates. J Bacteriol. 2005;187(17):6084–93. PubMed PMID: 1196134

24. Studier FW. Protein production by auto-induction in high density shaking cultures. Protein Expr Purif. 2005;41(1):207–34. PubMed PMID: 15915565.

25. Fraser CM, Norris SJ, Weinstock GM, White O, Sutton GG, Dodson R, et al. Complete genome sequence of *Treponema pallidum*, the syphilis spirochete. Science. 1998;281(5375):375–88. PubMed PMID: 9665876

26. Nielsen H. Predicting Secretory Proteins with SignalP. Methods Mol Biol. 2017;1611:59–73. Epub 2017/04/30. doi: 10.1007/978-1-4939-7015-5_6. PubMed PMID: 28451972.

27. Hiller K, Grote A, Scheer M, Münch R, Jahn D. PrediSi: prediction of signal peptides and their cleavage positions. Nucleic Acids Res. 2004;32(Web Server issue):W375–9. Epub 2004/06/25. doi: 10.1093/nar/gkh378. PubMed PMID: 15215414.

28. Juncker AS, Willenbrock H, Von Heijne G, Brunak S, Nielsen H, Krogh A. Prediction of lipoprotein signal peptides in Gram-negative bacteria. Protein Sci. 2003;12(8):1652–62. Epub 2003/07/24. doi: 10.1110/ps.0303703. PubMed PMID: 12876315.

29. Yu CS, Lin CJ, Hwang JK. Predicting subcellular localization of proteins for Gram-negative bacteria by support vector machines based on n-peptide compositions. Protein Sci. 2004;13(5):1402–6. Epub 2004/04/21. doi: 10.1110/ps.03479604. PubMed PMID: 15096640.

30. Yu NY, Wagner JR, Laird MR, Melli G, Rey S, Lo R, et al. PSORTb 3.0: improved protein subcellular localization prediction with refined localization subcategories and predictive capabilities for all prokaryotes. Bioinformatics. 2010;26(13):1608–15. Epub 2010/05/18. doi: 10.1093/bioinformatics/btq249. PubMed PMID: 20472543.

31. Berven FS, Flikka K, Jensen HB, Eidhammer I. BOMP: a program to predict integral beta-barrel outer membrane proteins encoded within genomes of Gram-negative bacteria. Nucleic Acids Res. 2004;32(Web Server issue):W394–9. Epub 2004/06/25. doi: 10.1093/nar/gkh351. PubMed PMID: 15215418.

32. Söding J, Biegert A, Lupas AN. The HHpred interactive server for protein homology detection and structure prediction. Nucleic Acids Res. 2005;33(Web Server issue):W244–8. Epub 2005/06/28. doi: 10.1093/nar/gki408. PubMed PMID: 15980461.

33. Bagos PG, Liakopoulos TD, Spyropoulos IC, Hamodrakas SJ. PRED-TMBB: a web server for predicting the topology of beta-barrel outer membrane proteins. Nucleic Acids Res. 2004;32(Web Server issue):W400–4. Epub 2004/06/25. doi: 10.1093/nar/gkh417. PubMed PMID: 15215419.

34. Kelley LA, Mezulis S, Yates CM, Wass MN, Sternberg MJ. The Phyre2 web portal for protein modeling, prediction and analysis. Nat Protoc. 2015;10(6):845–58. Epub 2015/05/08. doi: 10.1038/nprot.2015.053. PubMed PMID: 25950237.

35. Yang J, Yan R, Roy A, Xu D, Poisson J, Zhang Y. The I-TASSER Suite: protein structure and function prediction. Nat Methods. 2015;12(1):7–8. Epub 2014/12/31. doi: 10.1038/nmeth.3213. PubMed PMID: 25549265.

36. Wu S, Zhang Y. LOMETS: a local meta-threading-server for protein structure prediction. Nucleic Acids Res. 2007;35(10):3375–82. Epub 2007/05/05. doi: 10.1093/nar/gkm251. PubMed PMID: 17478507.

37. Giacani L, Molini B, Godornes C, Barrett L, Van Voorhis WC, Centurion-Lara A, et al. Quantitative analysis of *tpr* gene expression in *Treponema pallidum* isolates: differences among isolates and correlation with T-cell responsiveness in experimental syphilis. Infect Immun. 2007;75(1):104–12. PubMed PMID: 1828388

38. Cox DL, Luthra A, Dunham-Ems S, Desrosiers DC, Salazar JC, Caimano MJ, et al. Surface immunolabeling and consensus computational framework to identify candidate rare outer membrane proteins of *Treponema pallidum*. Infect Immun. 2010;78(12):5178–94. PubMed PMID: 20876295.

39. Centurion-Lara A, Giacani L, Godornes C, Molini BJ, Brinck Reid T, Lukehart SA. Fine Analysis of Genetic Diversity of the *tpr* Gene Family among Treponemal Species, Subspecies and Strains. PLoS Negl Trop Dis. 2013;16(7):e2222. PubMed PMID: 3656149

40. LaFond RE, Molini BJ, Van Voorhis WC, Lukehart SA. Antigenic variation of TprK V regions abrogates specific antibody binding in syphilis. Infect Immun. 2006;74(11):6244–51. PubMed PMID: 16923793.

41. Giacani L, Molini BJ, Kim EY, Godornes BC, Leader BT, Tantalo LC, et al. Antigenic variation in *Treponema pallidum*: TprK sequence diversity accumulates in response to immune pressure during experimental syphilis. J Immunol. 2010;184(7):3822–9. PubMed PMID: 20190145.

42. Reid TB, Molini BJ, Fernandez MC, Lukehart SA. Antigenic variation of TprK facilitates development of secondary syphilis. Infect Immun. 2014;82(12):4959–67. PubMed PMID: 25225245.

43. Salaun L, Snyder LA, Saunders NJ. Adaptation by phase variation in pathogenic bacteria. Adv Appl Microbiol. 2003;52:263–301. PubMed PMID: 12964248.

44. van der Woude MW, Baumler AJ. Phase and antigenic variation in bacteria. Clin Microbiol Rev. 2004;17(3):581–611. PubMed PMID: 15258095.

45. Saunders NJ. Evasion of antibody responses: bacterial phase variation. In: Oyston BHPCF, editor. Bacterial Evasion of Host Immune Responses. Cambridge: Cambridge University Press; 2003. p. 103–24.

46. Bulach DM, Seemann T, Zuerner RL, Adler B. The Organization of Leptospira at a Genomic Level. In: Chan VL, Sherman PM, Bourke B, editors. Bacterial Genomes and Infectious Diseases. Totowa, NJ: Humana Press; 2006. p. 109–23.

47. Edwards MJ, White GF, Butt JN, Richardson DJ, Clarke TA. The Crystal Structure of a Biological Insulated Transmembrane Molecular Wire. Cell. 2020;181(3):665–73.e10. Epub 2020/04/15. doi: 10.1016/j.cell.2020.03.032. PubMed PMID: 32289252.

48. Lauber F, Deme JC, Lea SM, Berks BC. Type 9 secretion system structures reveal a new protein transport mechanism. Nature. 2018;564(7734):77–82. Epub 2018/11/09. doi: 10.1038/s41586-018-0693-y. PubMed PMID: 30405243.

49. Hagn F, Etzkorn M, Raschle T, Wagner G. Optimized phospholipid bilayer nanodiscs facilitate high-resolution structure determination of membrane proteins. J Am Chem Soc. 2013;135(5):1919–25. Epub 2013/01/09. doi: 10.1021/ja310901f. PubMed PMID: 23294159.

